# Universal phase behaviors of intracellular lipid droplets

**DOI:** 10.1101/741264

**Authors:** Shunsuke F. Shimobayashi, Yuki Ohsaki

## Abstract

Lipid droplets are cytoplasmic micro-scale organelles involved in energy homeostasis and handling of cellular lipids and proteins. The core structure is mainly composed of two kinds of neutral lipids, triglycerides and cholesteryl esters, which are coated by a phospholipid monolayer and proteins. Despite the liquid crystalline nature of cholesteryl esters, the connection between the lipid composition and physical states is poorly understood. Here, we present the first universal intracellular phase diagram of lipid droplets, semi-quantitatively consistent with the in vitro phase diagram, and reveal that cholesterol esters cause the liquid-liquid crystal phase transition under near-physiological conditions. The internal molecules of the liquid crystallized lipid droplets are aligned radially. We moreover combine in vivo and in vitro studies, together with the theory of confined liquid crystals, to suggest that the radial molecular alignments in intracellular lipid droplets are caused by an anchoring force at the droplet surface. Our findings on the phase transition of lipid droplets and resulting molecular organization contribute to a better understanding of their biological functions and diseases.

---

Cholesterol is an essential component of cell membranes and its dysregulation leads to lethal diseases, such as arteriosclerosis and neurodegenerative disorders [1, 2]. Cholesterol is inserted into cell membranes owing to its hydrophobicity upon uptake and modulates the physical interaction between phospholipids in cell membranes, which can cause liquid-liquid phase separation of a homogeneous lipid membrane into liquid-disordered (L_d_) and liquid-ordered (L_o_) phases [3–6]. Some cholesterol is esterified and transported to lipid droplets (LDs), where it is stored as cholesteryl esters (CEs) [7]. CEs are known to be cholesteric liquid crystals and undergo liquid-liquid crystal phase transition as a function of system parameters (ex. temperature) [8]. Thus, CEs have the potential to induce the phase transition of LDs; however, the biophysics on the phase transition is poorly understood [9, 10].

To understand the liquid-liquid crystal phase (L-LC) transition in living cells, we cultured Huh7 cells (human hepatocarcinoma) with 170 *μ*M cholesterol for 24h. The methyl-*β*-cyclodextrin (M*β*CD) was used to dissolve hydrophobic cholesterol in hydrophilic culture medium (see the Method section). We observed the L-LC phase transition using polarized microscopy, which has been used to visualize liquid crystal phase [8], under near-physiological conditions while decreasing temperature (Fig. 1a). The temperature was controlled with a custom-made copper plate and thermoplate, and probed by a thermocouple inserted into a dummy chamber filled with water. The phase transition temperature was different among LDs in each cell, which may be attributed to the mass ratio difference between CEs and trygricerides (TGs). Interestingly, many of the cells cultured with 170 *μ*M cholesterol showed the coexistent states of liquid and liquid crystalline in a single LD at the intermediate temperature range (Fig. 1a). To statistically quantify the phase transition behavior, we counted as a function of temperature the probability that at least one lipid droplet in a single cell is at liquid crystalline state, *p*, with Huh7 cells (n=25). Interestingly, the probability monotonically decreases as temperature increases for the Huh7 cells cultured with 25, 50, and 170 *μ*M cholesterol (Fig. 1c); but, the slope is dependent on the added cholesterol concentration. At *T* = 40°C, almost all LDs were at L-phase (*p* ≈ 0); but, at *T* = 10°C, *p* = 16, 80, and 100 % for 25, 50, and 170 *μ*M cholesterol, respectively, indicating a clear correlation between the value of *p* and added cholesterol concentration. No LC LDs were observed even at *T* = 10°C for the cells cultured with 400 *μ*M oleic acid (OA) or only 10% FBS for 24h.

**FIG. 1.**
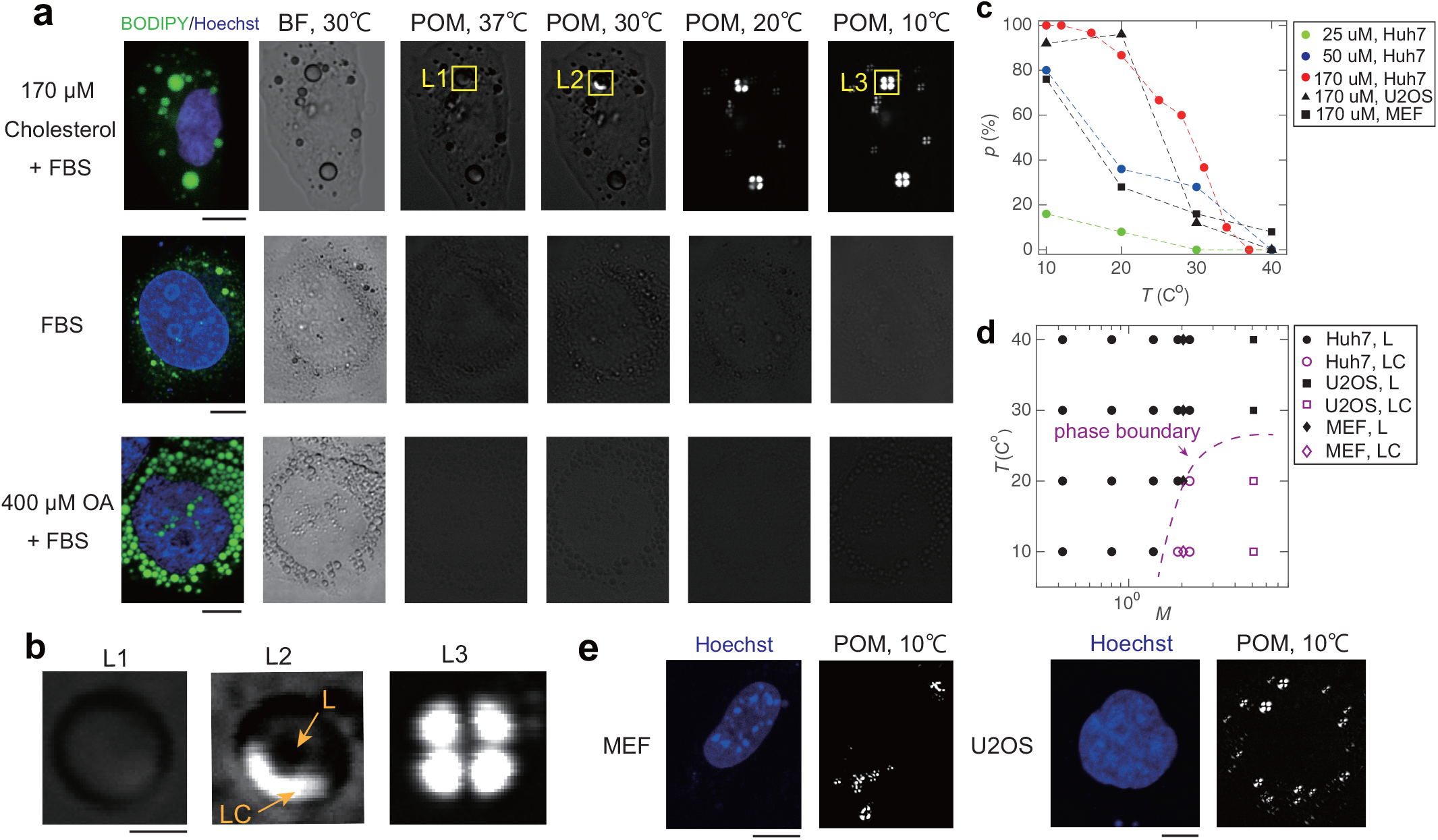
(a) Optical images of live Huh7 cells for three different of conditions (170 *μ*M cholesterol, only 10% FBS, and 400 *μ*M oleic acid). Bodipy and hoechst are markers of LDs and nucleus, respectively. The cross-polarized light was used for POM. Scale bars, 10 *μ*m. (b) Phase transitions in a LD. From the left, liquid (L) phase, the coexisting phase of liquid (L) and liquid-crystal (LC) phase, and liquid-crystal (LC) phase. Scale bars, 2 *μ*m. (c) Probability that at least one lipid droplet is at liquid crystalline state, *p*, as a function of temperature, *T*, for various cell types and cultured conditions. *n* = 25. (d) Universal intracellular phase diagram as functions of *T* and cholesterol ester/triglycerides (CE/TG) mass ratio, *M*. The phase was determined as L-phase if 0 ≦ *p* ≦ 50 and LC-phase if 50 < *p* ≦ 100. (e) Optical images of live MEF and U2OS cells cultured with 170 *μ*M cholesterol for 24h.

Based on the statistical data, we next mapped an intracellular phase diagram as functions of temperature and the CE/TG mass ratio, *M*, (Fig. 1b). To determine the CE/TG mass ratio, LDs were purified from cells and then the mass was measured by a TLC assay [11–13] (Fig. S1). The phase was determined to be L-phase if 0 ≦ *p* ≦ 50 and LC-phase if 50 < *p* ≦ 100. The phase diagram shows that LDs undergo the L-LC phase transition at the saturation mass ratio, *M*\*(*T*), with a fixed temperature and that *M*\*(*T*) decreases with a decrease in temperature. These findings suggest that CEs drive liquid crystallization of LDs.

We next investigated whether the phase transition phenomena is universal for other types of cells. We cultured MEF (mouse embryonic fibroblasts) and U2OS (human osteosarcoma) cells with 0.4 mM OA or only 10% FBS or 170 *μ*M cholesterol for 24h as we did with Huh7 cells. We never observed LDs with LC-phase for cells cultured with 0.4 mM OA or only 10% FBS; but, we observed LDs with LC-phase for MEF and U2OS cells cultured with 170 *μ*M cholesterol as well as with Huh7 cells (Fig. 1e). These behaviors are consistent with the behaviors in Huh7 cells (fig. 1c). The quantification of the CE/TG mass ratio for MEF and U2OS cells cultured with 170 *μ*M cholesterol and overlay plot of the data points in intracellular phase diagram give us the universal phase boundary of L-LC phase transition in living cells, independent of cell types (fig.1d). The universal intracellular phase diagram confirms that CEs work as drivers of liquid crystallization of LDs.

To get quantitative insights into how CEs induce liquid crystal transition in LDs, we reconstituted artificial liquid droplets (ALDs) in vitro systems and studied the phase behavior (Fig. 2a). ALDs were reconstituted with cholesteryl oleates (COs) and/or trioleins (TOs), which are one type of typical CEs and TGs, respectively, and MilliQ (18.2 MΩ cm), to which the surfactant (DOPC or Brij58 or Tween20) was dissolved with 3wt%. The surfactant solution was poured into COs which was pre-heated to *T* = 70°C to liquidify, and it was immediately pippeted for emulsification and cooled down to room temperature (*T* = 25 ± 1°C). This resulted in size-poilydisperse ALDs with *R* < 200 *μ*m, where *R* is the droplet diameter. Then, we mapped out the phase diagram of ALDs as functions of temperature *T* and the mole fraction of COs, *ϕ*_co_. Note that *ϕ*_co_ = 0.5 means the equal mole amounts of COs and TOs. We here focused on the ALDs with 1 ≤ *R* ≤ 70 *μ*m and did not observe any size-dependent structure change; also, the phase transition temperature is independent of *R* and the type of surfactants (Fig. S1). When the droplets were imaged under a polarized light microscope in the crossed Nicols condition, we observed three types of phases, i.e., L-phase, L-LC coexsiting phase, and LC-phase (Fig. 2b). The droplets with *ϕ*_co_ = 1 show L-LC phase transition at *T* = 38 ± 1 °C. Observing the transition dynamics, the nucleation of LC-phase randomly occurs, implying homogeneous nucleation (Fig. 2c, Movie S1). Then, the molecules start radially aligning from the droplet surface, resulting in a transition from polydomain pattern to the pattern of the Maltese cross (Fig. 2c). On the other hand, the droplets with *ϕ*_co_ = 0 are kept at the liquid state even at *T* = 0 ± 1°C. Interestingly, the droplets with 0.2 ≤ *ϕ*_co_ ≤ 0.8 exhibit the L-LC coexistent phase. Observing the dynamics, the randomly nucleated LC droplets grow with a power law of ~1/3 while diffusing in a droplet, suggesting diffusion-limited Ostwald ripening (Movie S2) [14]. We moreover realized that if the x-axis of the phase diagram, i.e., *ϕ*_co_, is changed to the mass ratio using the molecular weights of COs and TOs (651.1 and 885.4 g/mol, respectively), interestingly, the region we investigated in living cells (Fig. 1d) corresponds to the boxed region by the dashed-line (Fig. 2b). We then found that the phase boundaries of both phase diagrams are semi-quantitatively consistent (Fig. 1d, 2b). These findings suggest that LDs are two-component system of CEs and TGs, and CEs drive the L-LC phase transition.

**FIG. 2.**
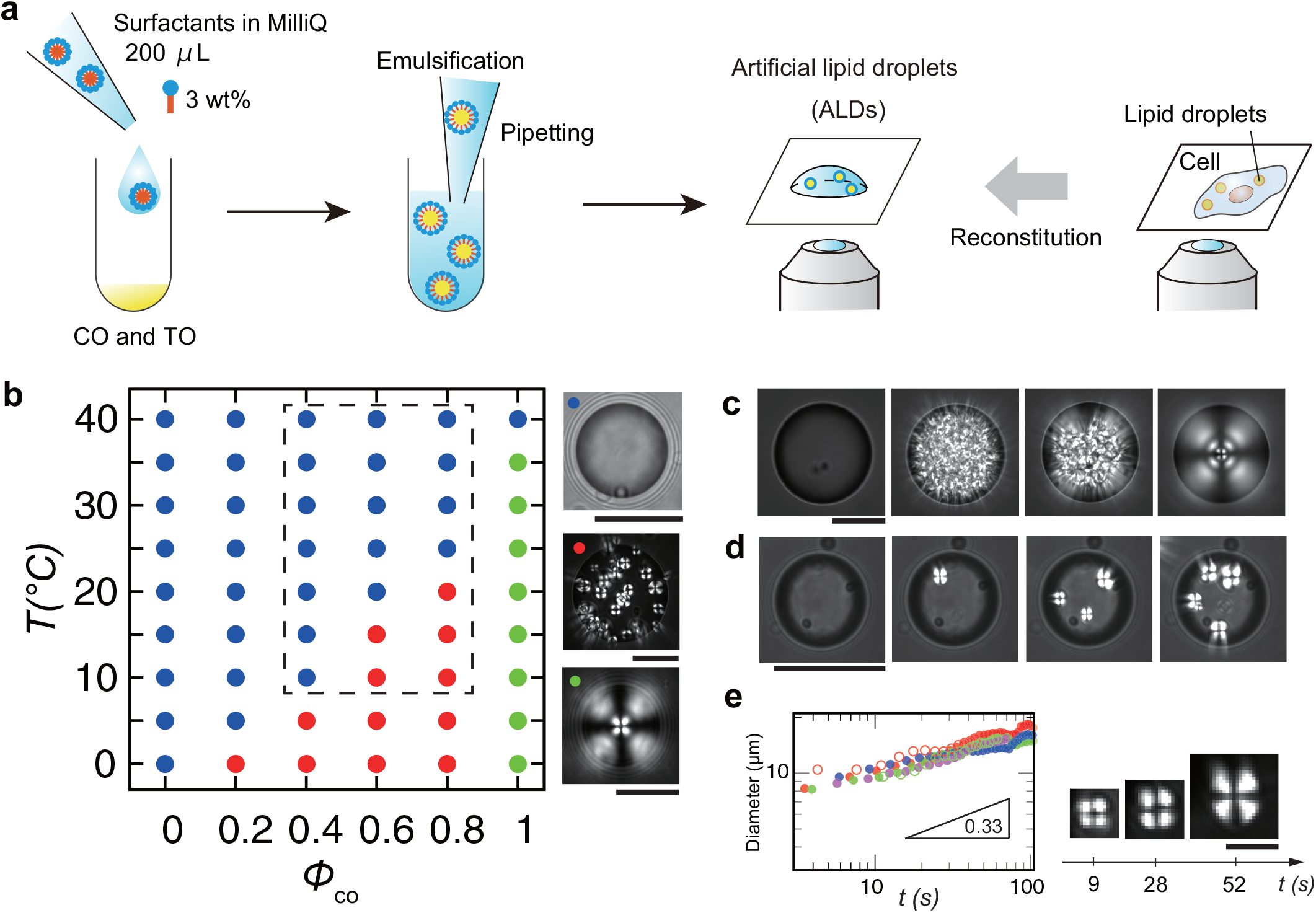
Phase behavior of artificial lipid droplets (ALDs). (a) Schematic diagram of the experimental set-up. (b) Phase diagram as functions of temperature *T* and mole fraction of cholesteryl oleates (COs), *ϕ*_co_. The blue, green and red circles represent the isotropic (L) phase, liquid-crystal (LC) phase, the coexistent phase of liquid and liquid-crystal (L-LC). On the right, the typical images of three phases are presented. The boxed region by the dashed-line corresponds to the investigated region in Fig.1 d. Scale bars, 100 *μ*m. (c) Dynamics of the L-LC phase transition of an ALD with *ϕ*_co_ = 1. Scale bar, 100 *μ*m. (d) Dynamics of the L-LC phase transition of an ALD with *ϕ*_co_ = 0.6. The LC-phase droplets nucleate and grow. Scale bar, 100 *μ*m. (e) Growth of nucleated LC droplets. Different colors or markers represent different droplets (left). Time-sequence of a LC droplet (right). Scale bar, 10 *μ*m.

We next sought to understand why the molecules are radially aligned both in intracelluar LDs and ALDs with *ϕ*_co_ = 1 and *R* ≤ 70 *μ*m below a certain phase transition temperature (Fig. 1d,2b). Interestingly, we found that the ALDs with *ϕ*_co_ = 1 and *R* > 70 *μ*m did not show the radial alignments but polydomain structure (Fig. 3a,b). To get insights into the biophysics behind the radial alignment observed in the intracellular LDs (Fig. 1b), we studied why the size-dependent structure transition occurs. We first mapped out a phase diagram of the structure as functions of temperature *T* and diameter *R* (Fig. 3b). We increased the temperature to *T* =47 ± 1 °C above the L-LC phase transition temperature *T_c_* (= 38 ± 1°C). All the droplets above *T_c_* were isotropic. The temperature was probed with an accuracy of ± 1.0°C using a thermocouple. Then, the temperature was decreased at a rate of −0.4°C/s to each point in the phase diagram. After about 10 min, the droplets were determined to be either in the radial-LC or polydomain-LC phase by means of cross-polarized light. The phase diagram clearly shows the structure transition between the radial and polydomain-LC phases in an all-or-none and size-dependent manner.

**FIG. 3.**
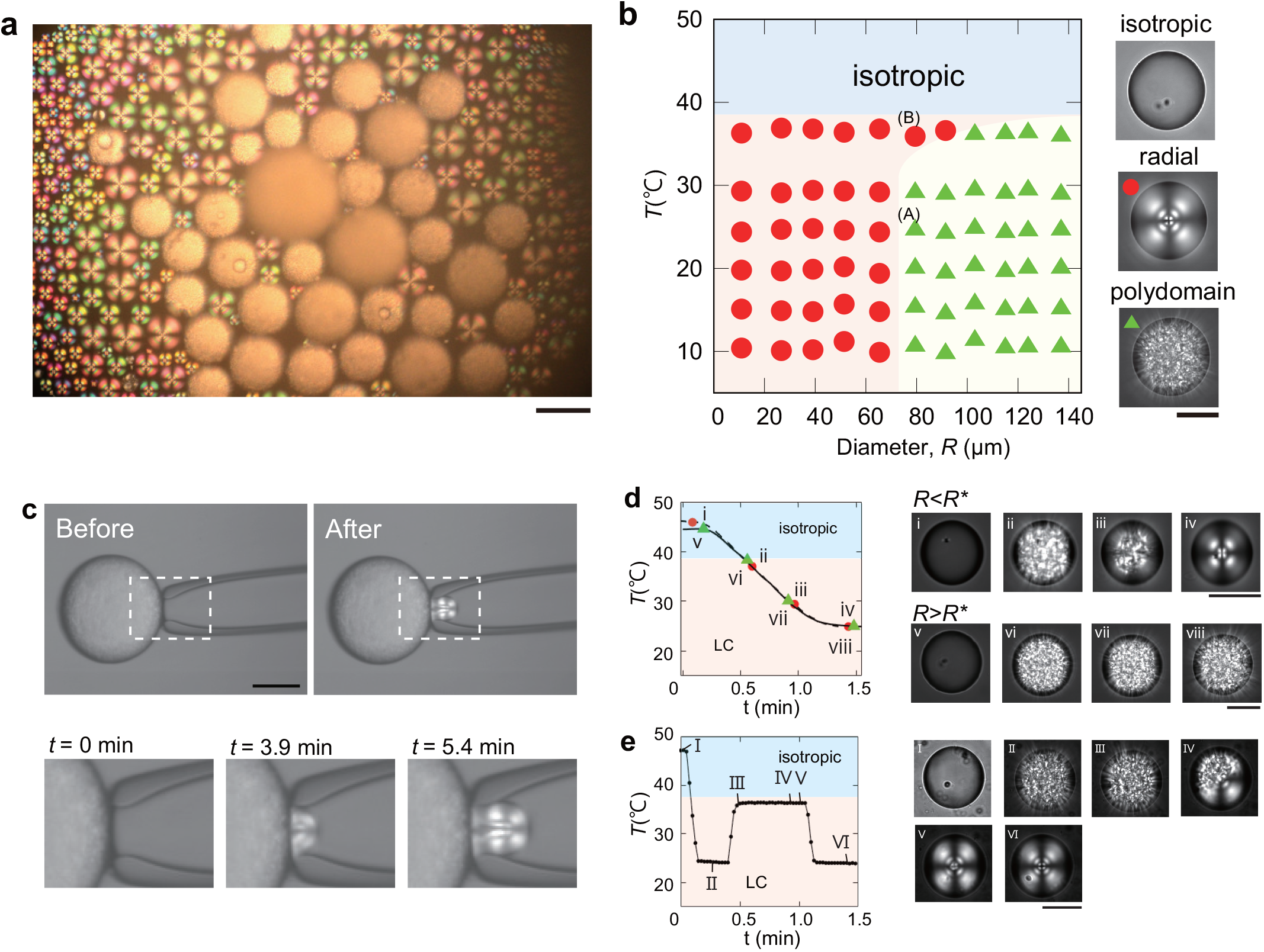
Size dependent structure transition of artificial lipid droplets (ALDs) with *ϕ*_co_ = 1. (a) An optical image of ALDs with crossed-polarizing light at *T* = 25 ± 1°C. The scale bar, 100 *μ*m. (b) Phase diagram as functions of temperature *T* and droplet diameter *R*. Red filled circles represent the radial-LC phase, where the molecules are radially aligned. Green filled triangles denote the polydomain-LC phase, where the molecules are aligned with a random direction in each domain. (c) Structure transition of ALDs with *ϕ*_co_ = 1 confined to a spherical cavity. (top) Polarized light microscopy images before (left) and after (right) micropipette aspiration are shown. (bottom) Sequential images (*t* = 0, 3.9, and 5.4 min, from left to right) of the rectangular region in top images under a constant pressure. Scale bars, 100 *μ*m. (d) Kinetic pathways of the isotropic-LC phase transition for two different sizes of droplets. (left) Temperature change as a function of time *t*. (right) Sequential polarized images for two pathways. Scale bars, 50 *μ*m. (e) Structure transition from the polydomain-LC to radial-LC phase during temperature cycle experiments. (left) Temperature change as a function of time *t* (right) Sequential polarized images. Scale bar, 50 *μ*m.

We next hypothesized from the phase diagram that the confinement of a part of the polydomain-LC droplet into a space with *R* < 70 *μ*m induces the transition to the radial-LC phase. We aspirated a droplet with the polydomain-LC phase using a micropipette (internal diameter, 60 *μ*m) connected with a microinjector at *T* = 25°C and pinched out a small droplet with *R* < *R*\* (Fig. 3c). Intriguingly, we found that the pinched drop with the radial-LC phase appears in the micropippet while the unpinched one remained the polydomain-LC phase (Movie 3). This finding suggests that the confinement to a spherical cavity with *R* < *R*\* gives rise to the structure change to radial-LC phase. This is another piece of evidence for the size-dependent configuration transition.

To get physical insights into why the molecules in the intracellular LC droplets are radially aligned, we consider the physical mechanism behind the size-dependent structure transition by means of the theory of confined liquid crystals. Considering cholesteryl oleates are in the smectic-A phase below *T* ≈38°C [15], total free energy of the confined smectic-A phase is expressed as [8, 16–19]

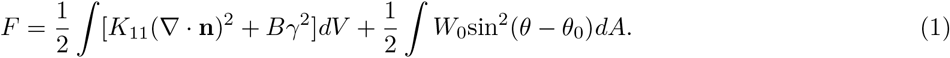

The first term in Eq.(1) is the contribution of splay deformations, and *K*_11_ denotes the splay Frank elastic constant and controls the splay of the layer normal, where n is the local nematic director. The terms of twist (*K*_22_(**n** · ∇ × **n**)^2^) and bend (*K*_33_(**n** × ∇ × **n**)^2^) deformations considered in the free energy of the nematic phase are not present due to the layer constraint [17, 18, 20]. The second term is the energy associated with layer dilation, *B* is compression modulus, and *γ* ≡ (*d* − *d*_0_)/*d*_0_ is a relative change in layer thickness from its equilibrium value *d*_0_ [17–19]. The ratio of splay and Young constants determines a length scale 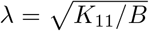, which is close to smectic layer thickness. When the typical radius of curvature of the distorted SmA is much larger than 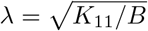, the dilation term is decoupled from the curvature term and dropped from the equation [18]. We therefore ignore this term. The third term is the surface anchoring energy, and *θ*_0_ and *θ* are the preferred and actual anchoring angles, respectively. In our system, it is assumed that the anchoring energy is due to the physical interaction between cholesterol esters and phospholipids at the surface. Finally, our free energy is reduced to 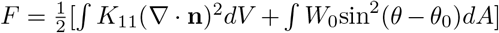, meaning that the free energy is the sum of the elastic energy in the droplet and anchoring energy at the surface.

Let us now compare free energies of the droplets with *R* between the radial-SmA phase and polydomain-SmA phase (denoted by *F*_rad_(*R*) and *F*_poly_(*R*), respectively). The gradient of the local nematic director, **∇** · **n**, in the radial-SmA phase is given by ~ 1/*R* and total integrated Frank energy *F*_el,rad_(*R*) is (*K*_11_/*R*^2^)*R*^3^ ~ *K*_11_*R*. On the other hand, **∇** · **n** in the polydomain-SmA phase is ~ 1/*ξ*, where *ξ* is the correlation length of local nematic alignment [21]. The elastic energy of each domain is ~ *K*_11_*ξ*, and total integrated energy *F*_el,poly_(*R*) is ~ *K*_11_*ξ*^−2^*R*^3^. Assuming that *R* ≫ *ξ*, the relation *F*_el,rad_(*R*) ≪ *F*_el,poly(*R*)_ is obtained. Moreover, the droplet is coated by the amphiphile surfactant. The hydrophobic part that faces toward the droplet center gives rises to the strong homeotropic anchoring on the nematic director at the droplet surface [22]. This situation can lead to *F*_anc,rad_ ≤ *F*_anc,poly_, where *F*_anc,rad_ and F_anc poly_ are the anchoring energies for the radial-SmA and polydomain-SmA phases, respectively. Finally, the relation *F*_rad_(= *F*_el,rad_ + *F*_anc,rad_) ≪ *F*_poly_(= *F*_el,poly_ + *F*_anc,poly_) is obtained, suggesting that the polydomain Sm-phase is energetically unstable, independent of the droplet size. Observing the two kinetic pathways of the L-LC phase transition of droplets above or below *R*\*, the polydomain structure emerges on both pathways immediately after *T* passed through *T*_c_ (Fig. 3d, Movie S4,S5). Then, the radial alignment starts only for the droplets with *R* < *R*\*. We therefore conclude that the polydomain phase is a kinetically trapped metastable state.

Moreover, we experimentally investigated whether the polydomain-SmA phase is metastable. We first prepared the droplet with *R* ≈ 80 *μ*m in the isotropic phase at *T* = 47 ± 1 °C and decreased the temperature to *T* = 24 ± 1 °C at a rate of −0.4°C/s (Fig. 3b-A), resulting in emergence of the polydomain-SmA phase (Fig. 3e, Movie S6). The temperature was then increased to *T* = 36 ± 1 °C at the same rate (Fig. 3b-B) while the phase boundary was crossed between the two phases. Immediately, the radial alignment started from the droplet interface, leading to transition to the radial-SmA phase (Fig. 3e). Even if the temperature was returned to *T* = 25 ± 1°C, the radial-SmA phase remained stable. These results provide evidence that the polydomain-SmA phase is the metastable state.

We now consider why the metastable polydomain Sm-phase undergoes the transition to the radial-Sm phase only for the droplets with *R* < *R*\*. Looking at the free-energy functional of the LC droplets (Eq.(1)), the driving force to overcome the volumetric elastic energy should come from the surface anchoring. The free energy gain due to the favorable surface anchoring corresponds to ~ *W*_0_*R*^2^ and the structure transition to the radial-SmA phase should take place when the energy gain ~ *W*_0_*R*^2^ overcomes the elastic energy *F*_el,poly_ ~ *K*_11_*ξ*^−2^*R*^3^. Therefore, the threshold droplet diameter *R*\* is obtained as

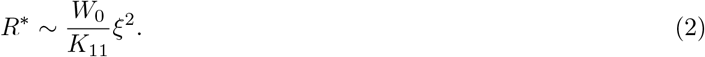

Besides, can be written as 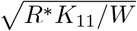 and estimated to be on the order of 1 *μ*m with typical values for *K*_11_ ~ 10^−11^ N [17, 22, 23], *W*_0_ ~ 10^−4^ J/m^2^ for the “strong anchoring” [16, 24], and *R*\* ≈ 70 *μ*m. The *ξ*-value is comparable with the value observed in liquid crystalline polymers and elastomers [21] and is consistent with the optical observations (Fig. 3b). We conclude that the radial alignments for the droplets with *R* < *R*\* are driven by the homeotropic anchoring force at the droplet surface and suggest that the radial alignments in LDs in living cells are caused by the anchoring force at the droplet surface (Fig. 1).

We also found that *R*\* remained constant to be at ~70 *μ*m below *T* ≲ 30 °C; on the other hand, *R*\* increased as *T* approached *T_c_* and reached approximately 100 *μ*m at *T* = 36 ± 1°C (Fig.3b). We theoretically consider the mechanism behind the temperature dependence of *R*\*. As depicted in Eq.(2), *R*\* is determined by the balance between *W*_0_, *K*_11_ and *ξ*. Elastic constant *K*_11_ has a second order in the nematic order parameter *Q* and *W*_0_ is linear power of *Q* [22], leading to *W*_0_/*K*_11_ ~ *Q*^−1^. On the other hand, *ξ* experimentally seems to be constant so that the temperature dependence of *ξ* is likely minor in our experiments. Therefore, because *Q* → 0 as *T* – *T_c_, R*\* = *W*_0_/*K*_11_ diverges as *T* → *T_c_*. In other words, the relative contribution of the surface alignment increases as *T* → *T_c_*. Also, *R*\* remained constant below *T* ~ 30 °C; this would be attributed to the near-saturation of the *Q* value when *T* ≪ *T_c_* [25].

There is a mounting evidence that phase transitions play important roles in living cells, particularly associating with membrane-less organelles or condensates [3, 14, 26–28]; nevertheless, there has been little work on the phase transition in lipid droplets [29]. Our study to quantify the universal intracellular phase diagram, semi-quantitatively consistent with the in vitro phase diagram, is only the beginning of the fundamental biophysics for lipid droplets. We moreover combine in vivo and in vitro studies, together with the theory of confined liquid crystals, to suggest that the radial molecular alignments in LDs in living cells are caused by the anchoring force at the droplet surface. However, it is still not clear whether the radial alignments cover the whole droplet or only near the LD surface, as discussed on LDs or LDL for several decades [9, 30–35]. The structural details must be revealed by the precise investigation using both polarized optical microscopy and cryo-EM in the future. Such a deeper understanding of the LD structure will contribute to biological functions and diseases on LDs. For example, macrophages that are highly loaded with CEs can cause the lipid ordering in LDs and differentiate to foam cells; these cells are the main cellular components of fatty streaks and indicate the earliest stage of atherosclerosis [9, 10]. Investigation of the relation between the structure change due to the liquid-liquid crystal phase transition and the onset of differentiation to foam cells might give us novel insights into the mechanism behind arteriosclerosis.

## MATERIAL AND METHODS

### The cell line

Huh7 cell lines were obtained from the Japanese Collection of Research Bioresources Cell Bank. MEF and U2OS were kindly donated by Dr. Noboru Mizushima (Tokyo University) and Dr. Hidemasa Goto (Aichi Cancer Center), respectively. Cells were cultured in MEM (for Huh7) or DMEM (for other cells) supplemented with 10% of fetal bovine serum (FBS) and antibiotics at 37°C in a humidified atmosphere of 95% air and 5% *CO*_2_. In some experiments, the cells were cultured at a certain concentration of OA (Sigma-Aldrich) in complex with fatty-acid-free bovine serum albumin (Wako) at a molar ratio of 6:1. In other experiments, the cells were cultured at a certain concentration of cholesterol (Sigma-Aldrich) in complex with methly-*β*-cyclodextrin (Sigma-Aldrich) at a molar ratio of 1:6.

### LD purification and TLC assay

LD fraction was purified by nitrogen cavitation and sucrose density ultracentrifugation as described before (Fujimoto T, 2001). Lipids were extracted (Bligh and Dyer, 1959) and separated on HPTLC Silica gel 60 (Merck) with a hexane-diethylether-acetic acid (80:20:1), and charred by 3% copper acetate in 85% phosphoric acid at 180C.

### Microscopy and imaging

Polarized light imaging was performed under an inverted microscope (Nikon Ti-E) with a confocal laser scanning system (Nikon A1), equipped with a ×4, 0.13 NA, ×10, 0.45 and ×20, 0.75 NA dry objective lens (Nikon, Japan). The images were captured by a Andor iXon3 EM-CCD camera or canon NY-X6i colored camera.

### Micromanipulation

This procedure was performed using an Narishige micromanipulator (MM-92) and pneumatic micro-injector (IM-11-2) mounted on an inverted microscope (Nikon Ti-E), equipped with a glass micropipette (60 *μ*m internal diameter, VacuTip FCH, Eppendorf).

### Temperature control and logging

Temperature was controlled by means of a custom-made copper plate with thickness 200 *μ*m mounted on a thermoplate (CHSQ-C, Tokai hit). The copper plate had a hole of 4 mm in diameter at the center. The temperature was probed by a thermocouple inserted into a dummy chamber filled with water and was logged.

